# Solution-state NMR Assignment and Secondary Structure Analysis of the Monomeric *Pseudomonas* Biofilm-Forming Functional Amyloid Accessory Protein FapA

**DOI:** 10.1101/2023.07.18.549541

**Authors:** Chang-Hyeock Byeon, Ümit Akbey

## Abstract

FapA is an accessory protein within the biofilm forming functional bacterial amyloid related fap-operon in *Pseudomonas*. We present a complete sequential assignment of ^1^H_amide_, ^13^C_α_, ^13^C_β_, and ^15^N NMR resonances for the functional form of the monomeric soluble FapA protein, comprising amino acids between 29-152. From these observed chemical shifts, the secondary structure propensities (SSPs) were determined. FapA predominantly adopts a random coil conformation, however, we also identified small propensities for α-helical and β-sheet conformations. Notably, these observed SSPs are smaller compared to the ones we recently observed for the monomeric soluble FapC protein. These NMR results will provide valuable insights into the activity of FapA in functional amyloid formation and regulation, that will also aid developing strategies targeting amyloid formation within biofilms and addressing chronic infections.

## Biological Context

Biofilm-forming functional bacterial amyloid (FuBA) fibrils differ from pathological amyloids due to their optimized evolutionary sequences, tight regulation, and vital *in vivo* roles. Their crucial participation in biofilm formation maintains the integrity of bacterial communities within biofilms, which are responsible for chronic human infections that pose challenges in treatment and contribute to antimicrobial resistance (AMR). Well-studied examples of FuBAs include CsgA, TasA, FapC, and PSMs.(Chapman, Robinson et al. 2002, Chiti and Dobson 2006, Dueholm, Petersen et al. 2010, Romero, Aguilar et al. 2010, Chiti and Dobson 2017, Akbey and Andreasen 2022) Targeting FuBAs within biofilms could introduce a novel approach to combat chronic infections, but the lack of structural information and understanding of *in vivo* regulation present significant challenges.

The fap-operon in *Pseudomonas* consists of six proteins, fapA-F. (Dueholm, Petersen et al. 2010) These proteins control the production of functional amyloids and their related accessory proteins, which are involved in the biogenesis of biofilm-forming amyloids, **Figure 1A**. In a recent study, we achieved a significant milestone by presenting a complete solution NMR resonance assignment of monomeric FapC, the major amyloid protein responsible for the biofilm formation in *Pseudomonas*.(Byeon, Wang et al. 2023) Building on this progress, we extend our NMR analysis to provide a complete solution-state NMR resonance assignment of a key accessory protein FapA within the fap-operon. Additionally, we discuss the NMR chemical shift-based secondary structure propensities of soluble FapA to enhance our understanding of its structural features. We have also recently shown that FapA is an important interaction partner of FapC and modifies its aggregation propensity.(Byeon, Hansen et al. 2023) We and others proposed that FapA has chaperone properties that slows down FapC fibrillation, a process which should be prevented.(Byeon, Hansen et al. 2023, Rasmussen, Kumar et al. 2023) Uncontrolled spontaneous fibrillation would be detrimental to bacteria; thus, FapA and other accessory proteins in the fap-operon tightly control fibrillation process until amyloids are secreted outside the cell. Elucidating the molecular mechanism of FapA, belonging to a unique class of proteins capable of acting as a chaperone or chaperone-like, holds significant potential for developing strategies to prevent amyloid formation in biofilms and addressing related chronic infections. Chaperones exhibit diverse effects on fibrilization, as seen in both pathological and functional amyloids, ranging from slowing down fibril formation to prevention or even reversal.(Ulamec, Brockwell et al. 2020) Notably, CsgE decelerates fibril formation of functional amyloid CsgA,(Sewell, Stylianou et al. 2020) while chaperones from the HSP90 family delay pathological αSynuclein amyloid fibrillation of.(Bohush and Filipek 2020)

**Figure 1.**
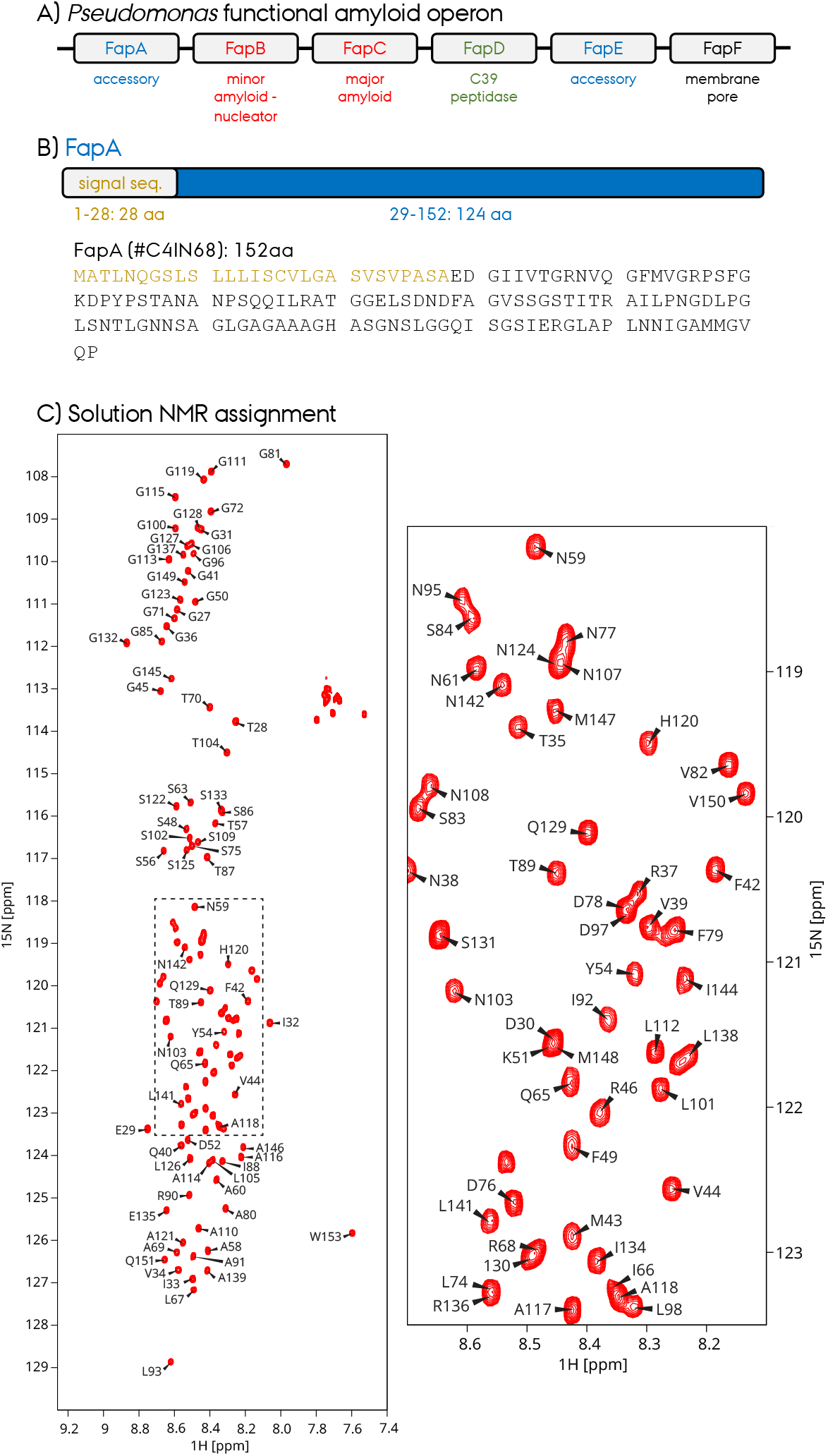
A) Biofilm forming functional amyloid related operon from *Pseudomonas*, the “fap-operon”. B) The domain organization and amino acid sequence of FapA. The signal sequence is between amino acids 1-28, whereas the functional secreted FapA is between amino acids 29-152. C) ^1^H-^15^N HSQC NMR spectra of FapA 29-152 with complete NMR resonance assignment. The section of the spectrum which had more densely packed peaks is shown as a zoomed-out region.

In this study, we present the solution-state NMR assignment of FapA, which comprises 152 amino acids (**Figure 1B**). Our investigation focuses on the functional *in vivo* form of FapA specifically amino acids between 29-152, without signal sequence (amino acids 1-28), which guides the protein to the secretion chamber and is cleaved upon secretion into the periplasm. Upon analyzing the 2D HSQC fingerprint NMR spectrum of isotope-labeled FapA (**Figure 1C**), it is apparent that FapA exhibits characteristics of an intrinsically disordered protein (IDP) with limited chemical shift dispersion, similar to FapC.(Byeon, Wang et al. 2023) Through the complete assignment of backbone NH resonances, we have observed that FapA predominantly adopts a disordered structure with measurable secondary structure propensities (SSPs) as α-helices and β-sheets. These SSPs are comparatively smaller than those observed in FapC. Our study provides essential NMR resonance assignments, enabling future investigations into the structural characteristics of soluble FapA and its interaction with e.g. FapC. A comprehensive understanding of chaperone activity of FapA, combined with its structural properties, will offer valuable insights in amyloid formation and regulation, as well as will aid the development of strategies targeting amyloid formation within biofilms and addressing chronic infections.

### Plasmid Construction

We followed the protocol we established recently for the expression of a short FapC construct. (Byeon, Wang et al. 2023) Recombinant FapA FL (residues 29-152, 124 amino acid) plasmid was cloned into pET32 vector using the In-fusion cloning mechanism (Takara Bio). The FapA construct was expressed with an N-term Thioredoxin (Trx), a 10 histidine (His10) tag, and a thrombin cleavage site. The final cleaved FapA protein contains five additional non-native amino acids, four at the N-terminus from cloning/cleavage artifacts (GSGT), and one at the C-terminus to increase UV280 absorbance for quality control purposes during purification and protein quantification, since FapA does not have any native tryptophan.

### Recombinant protein production

Bacterial plasmids were transformed into BL21(DE3) *E. coli* bacteria. All nonisotope-labeled samples were prepared in LB media. Isotope labeling for NMR studies was carried out by growing in minimal media with ^13^C-glucose (2g/L) and ^15^N-ammonium chloride (1g/L) as carbon and nitrogen sources, respectively. The transformed BL21(DE3) cells were plated onto LB agar containing ampicillin and grown overnight in 37 °C incubator. The colonies were resuspended in the corresponding media with ampicillin and then inoculated into the large volume. These cultures were grown in shaker incubator at 37 °C until the OD600 reached a value between 0.8-1.0 OD. IPTG was added to 1 mM final concentration to induce protein expression and grown for another 3-4 hours at 37 °C. Cells were harvested by centrifugation (7200 RCF, for 20 min at 4 °C). While on ice, the cells were lysed in lysis buffer (50 mM tris, 300 mM sodium chloride, 0.2 mM phenylmethylsulfonyl fluoride (PMSF), pH 8) using sonication with two cycles of 5 min at 70% power with 50% duty cycle (1 sec on and 1 sec off). The lysed cells were then centrifuged for 20 min at 4 °C and the soluble fraction was collected for further purification.

His-Tag affinity column was used with a gradient elution from 25 mM to 350 mM imidazole over 140 mL to initially purify Trx-FapA. The protein was then dialyzed into Dialysis buffer (20 mM tris, 0.02% sodium azide (w/v), pH 8.7). The sample was then processed with thrombin at 8U per mg of Trx-FapA protein at 4 °C for 12-16 hr. Complete processing was verified by SDS-PAGE and mass spectrometry, and then PMSF for a final concentration of 0.2 mM to stop the process. The cleaved FapA was further purified using SourceQ anionic exchange column with a linear gradient from 0 mM – 50 mM sodium chloride over 60 mL volume in Dialysis buffer with 0.2 mM PMSF. Proteins of ∼0.3 mM concentration in 20 mM Sodium Phosphate, 1 mM D6-DSS, 10% D_2_O, pH 7.8 buffer were made for the NMR experiments. The NMR samples were stored at 4°C before the NMR experiments.

### NMR Spectroscopy

Uniformly ^13^C,^15^N isotope labeled FapA FL (124 amino acid, without the signal peptide, Figure 1B) was used to record 2D ^1^H-^15^N HSQC and 3D HNCACB, HN(CO)CACB spectra. All spectra were acquired at Bruker Avance III 600 MHz spectrometers equipped with a 5 mm triple resonance TCI cryoprobes. 5 mm NMR sample tubes were used with a total volume of 550 μl sample and a protein concentration of ∼300 μM. The sample was stored at 4°C before performing NMR experiments, and the spectra were recorded at 274 K. **Table 1** shows the details of the NMR experiments. The spectra were processed by Topspin 3 (Bruker Biospin). The NMR resonance assignment was done manually by using CCPNmr Analysis 3.1.(Stevens, Fogh et al. 2011) The ^1^H chemical shifts were referenced to 0 ppm by using DSS as an internal standard added to the NMR samples. The ^13^C and ^15^N chemical shifts were indirectly referenced by using the ^1^H frequency as explained previously.(Wishart, Bigam et al. 1995)

**Table 1.** Parameters of the 2D and 3D NMR experiments performed on 300 μM ^13^C,^15^N FapA in this study. All spectra were recorded at 274 K and in 20 mM Sodium Phosphate, 1 mM D6-DSS, 10% D_2_O, and pH 7.8. Chemical shift calibration was done using the D_6_-DSS as an internal reference.

| NMR Experiment | ns | RD | TD direct | Increment | SW | TD | Increment | SW | TD | Increment | SW | Instrument | Expt. Time |
| --- | --- | --- | --- | --- | --- | --- | --- | --- | --- | --- | --- | --- | --- |
| | | s | 1H | $\mu\text{s}$ | ppm | 15N | $\mu\text{s}$ | ppm | 13C | $\mu\text{s}$ | ppm | MHz | hour |
| 2D HN HSQC | 32 | 1 | 2048 | 52 | 16 | 198 | 685.0 | 24 | - | - | - | 600 | 1 |
| 3D HNCACB | 64 | 0.9 | 2048 | 52 | 16 | 64 | 685.0 | 24 | 36 | 110.4 | 60 | 600 | 72 |
| 3D HN(CO)CACB | 64 | 0.9 | 2048 | 52 | 16 | 64 | 685.0 | 24 | 36 | 110.4 | 60 | 600 | 72 |

### NMR Assignment, Statistics and Data Deposition

The 2D ^1^H-^15^N HSQC spectrum of FapA 29-152 is shown in **Figure 1C** with the complete backbone NH resonance assignment excluding the eight prolines. The summary of overall assignment statistics is as follows for the 124 amino acid FapA: 100% backbone HN nuclei (116/116, due to 8 proline), 100% of the backbone C_α_ (124/124), 100% of the C_β_ (101/101, due to 23 glycines). All the chemical shifts have been deposited in BMRB (access number #52035). A representative sequential NMR assignment stretch is shown in **Figure 2** for residues S56-N61 sandwiched between two proline residues based on the 3D HNCACB spectra. Resonances from the i (residue itself) and i-1 (preceding residues) are labeled accordingly.

**Figure 2.**
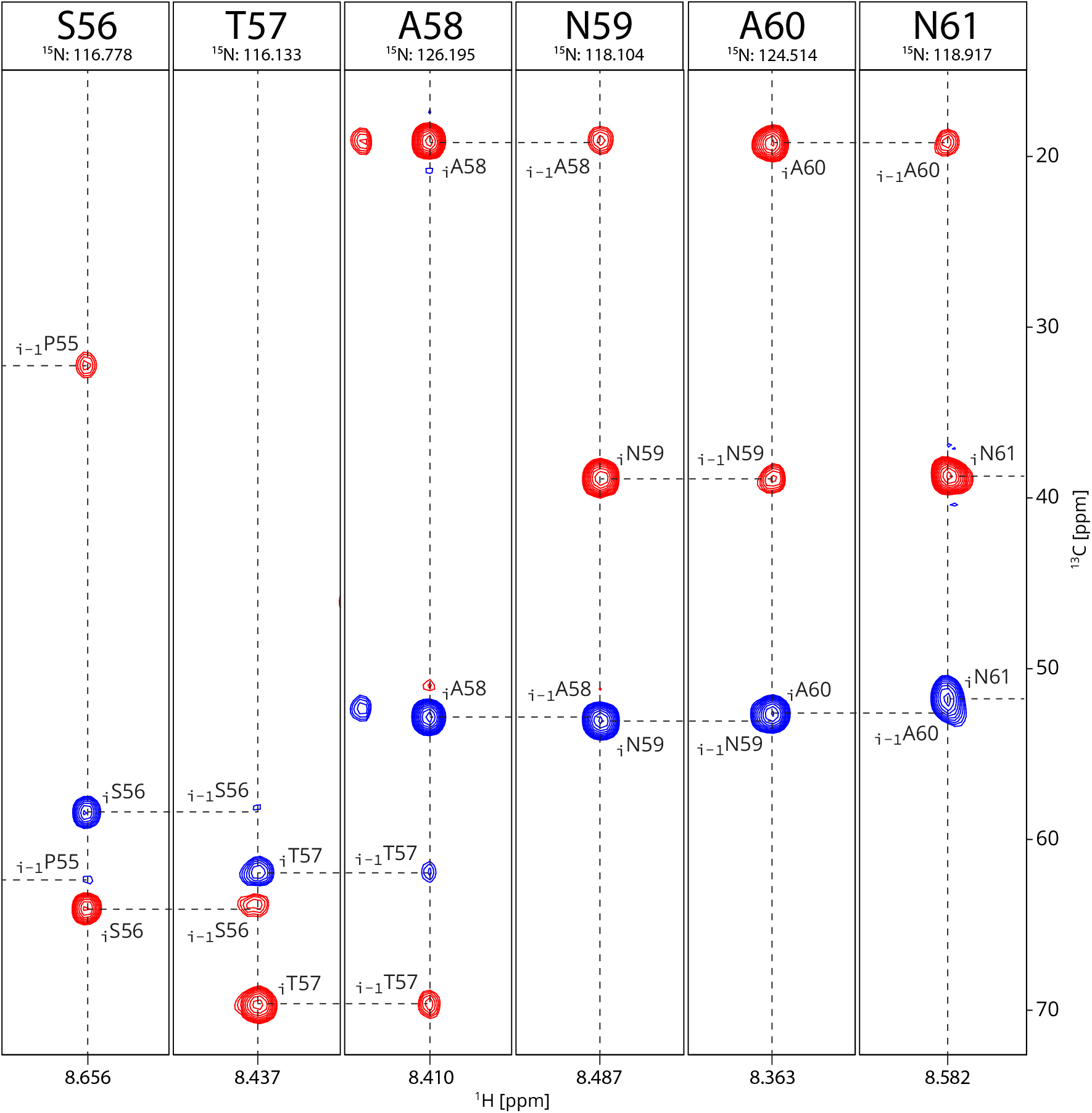
Representative sequential NMR resonance assignments for FapA for residues S56-N61. 3D HNCACB NMR spectrum shows the C_α_ resonances in blue and C_β_ resonances in red.

By using these C_α_, C_β_ NMR chemical shifts (ββ(C_α_)-ββ(C_β_)), we determined the deviation from random coil chemical shift values to identify stable α-helices or β-sheets. These residues have an average of 0.23 ppm deviation from random coil, data not shown here.(Wishart, Sykes et al. 1991) A majority of these values fall below the +/-1.4 ppm threshold and range between 1.4 ppm and -0.8 ppm. This indicates an IDP nature of the FapA with close to the random coil conformation and small structural propensities. With the full backbone assignment of FapA, the secondary structure analysis was performed. The neighbor corrected secondary structure propensity (ncSSP) method optimized for IDP proteins based on the work by Tamiola *et al*. was utilized to determine the SSPs from these chemical shifts, **Figure 3**. (Tamiola, Acar et al. 2010, Tamiola and Mulder 2012) ncSSP method corroborated that FapA has an IDP like nature. ncSSP analysis indicate a maximum SSP of ∼14.4%, which is smaller compared to our FapC result in which a maximum propensity of ∼25% was observed.(Byeon, Wang et al. 2023) 90 residues out of the 124 residues show α-helices SSPs, whereas 32 residues show β-sheet SSPs.

**Figure 3.**
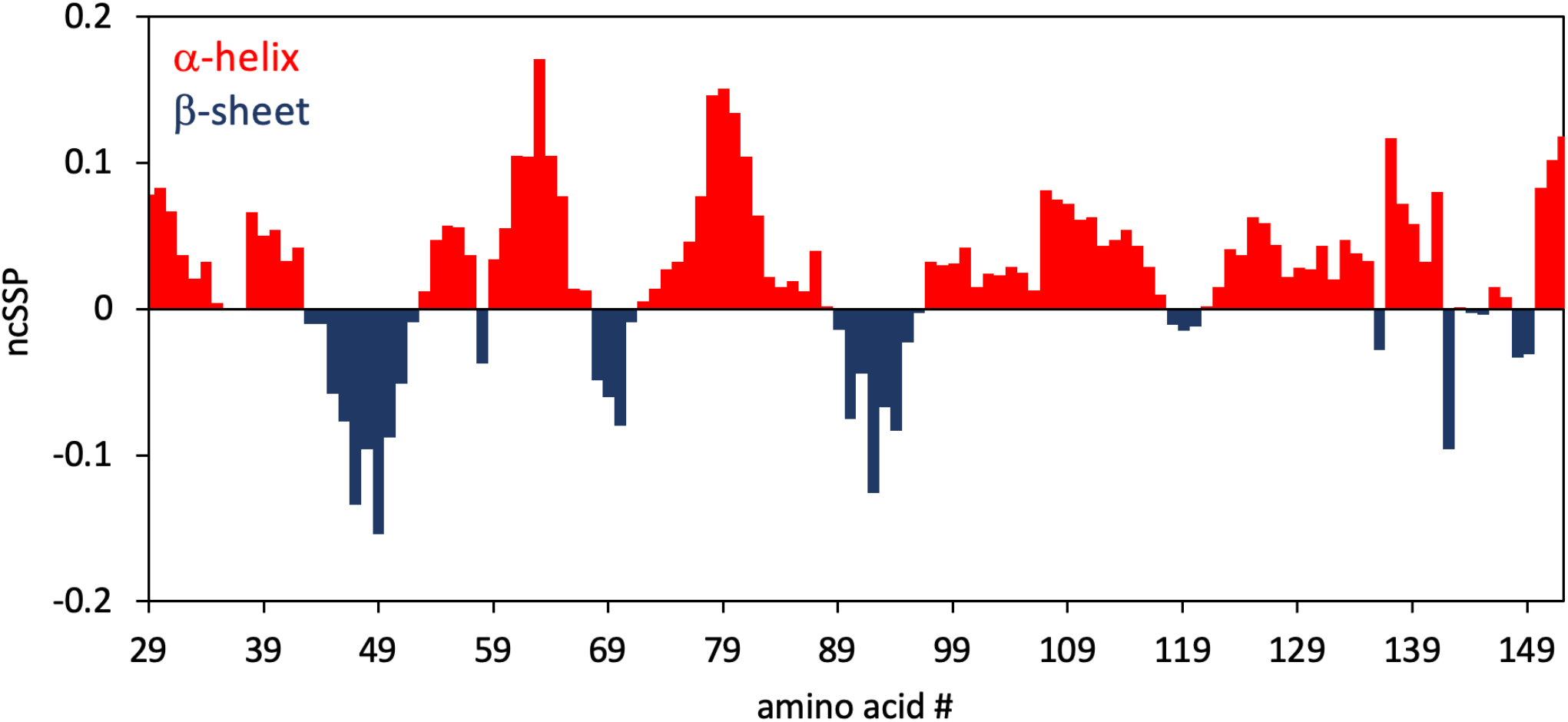
Residue-specific secondary structure propensities (red for α-helix and blue β-sheet) based on the NMR chemical shift via the neighbor-corrected secondary structure propensities (ncSSP) algorithm following the work by Tamiola *et al*.(Tamiola, Acar et al. 2010, Tamiola and Mulder 2012) The propensities are shown in terms of percentage, whereby 1 equal to 100% propensity.

### Summary

In summary, here we presented a complete sequential ^1^H_amide_, ^13^C_α_, ^13^C_β_, and ^15^N assignment of NMR resonances for the functional form of FapA protein. We also determined the secondary structure propensities (SSPs) from the observed chemical shifts. FapA adopts predominantly a random coil conformation. Nevertheless, small propensities for α-helical and β-sheet conformations were determined. These SSPs are at a smaller magnitude compared to the ones we observed for the FapC protein. These NMR resonance assignments will facilitate the structural studies of FapA and its interactions with other functional-amyloid related proteins from *Pseudomonas* or small molecules.

## Data Availability

NMR assignments been deposited in BMRB with access number #52035. Data will be available upon request.

## Author Contribution

Conceptualization: UA; Methodology: CHB, UA; Formal analysis and investigation: CHB, UA; Writing - original draft preparation: CHB, UA; Funding acquisition: UA; Resources: UA; Supervision: UA.

## Acknowledgement

We thank In-ja Byeon for helpful discussions and help in recording the 3D NMR spectra. Frans Mulder and Maria Andreasen are acknowledged for helpful discussions. UA acknowledges financial support from the CMRF Award from UPMC and University of Pittsburgh, University of Pittsburgh startup funding, and the high-field NMR infrastructure at the Structural Biology Department, School of Medicine, University of Pittsburgh.

